# MicroProphet: A Digital Twin Framework for Predicting Microbial Community Dynamics with Personalized Precision

**DOI:** 10.1101/2025.05.08.652793

**Authors:** Yuli Zhang, Kouyi Zhou, Xiaoke Chen, Haohong Zhang, Jin Han, Kang Ning

**Author notes:** **Corresponding authors** Further information and requests for resources should be directed to and will be fulfilled by the lead contact, Kang Ning. These authors contributed equally to this work (Yuli Zhang, Kouyi Zhou, Xiaoke Chen).

## Abstract

The ability to accurately predict the dynamic evolution of microbial communities is critical for advancing personalized medicine, precision intervention, and ecological system management. However, the irregular sampling, high missingness, and complex temporal behaviors that characterize longitudinal microbiome datasets present substantial challenges to reliable forecasting. Here we propose MicroProphet, a personalized digital twin framework capable of accurately forecasting microbial abundance trajectories from incomplete longitudinal observations without the need for data interpolation. By leveraging a time-aware Transformer architecture, MicroProphet reconstructs individualized microbial trajectories using as little as the initial 30% of time points, capturing critical transitional states through its attention mechanism. We demonstrate its robust cross-ecosystem generalizability across synthetic communities, human gut microbiomes, infant gut development, and corpse decomposition. In clinical contexts, MicroProphet enables early identification of disease-related microbial shifts and supports intervention timing optimization, exemplified in inflammatory bowel disease and antibiotic perturbation responses. By transforming incomplete and sparse data into actionable forecasts, MicroProphet establishes a foundation for real-time microbial monitoring, therapeutic decision support, and precision ecological management, paving the way for broader applications of digital twin systems in biology and personalized healthcare.

**Highlights:**

- MicroProphet establishes a personalized digital twin framework for forecasting biological dynamics from incomplete longitudinal observations, enabling precision health monitoring and intervention planning.
- By leveraging a transformer-based architecture, MicroProphet accurately reconstructs microbial community trajectories using as little as 30% of initial time points, without the need for data interpolation.
- The framework demonstrates robust cross-ecosystem generalizability across clinical, early-life development, and forensic contexts.
- MicroProphet empowers real-time tracking of microbial shifts, early detection of disease-associated transitions, and timing optimization for microbiome-targeted therapeutic strategies.

## Introduction

Microbial communities play crucial roles in maintaining the health of various ecosystems, ranging from the human-associated microbiota to agricultural and industrial environments.^1-3^ With the growing emphasis on precision medicine, the dynamic changes in microbial communities have become an important direction for understanding individual health. Unlike static, cross-sectional datasets that provide snapshots of microbial composition, longitudinal microbiome data reveal the evolving trajectories of microbial interactions and their ecological consequences, offering deeper mechanistic insights into community functions over time. However, the incompleteness and sparsity of such data pose substantial modeling challenges, limiting their translational potential.^4-6^

Classical modeling frameworks, such as generalized Lotka-Volterra systems (e.g., MDSINE),^7^ dynamic Bayesian networks (e.g., CGBayesNets),^8^ and digital twin models based on conditional inference trees (e.g., Q-net),^9^ have yielded important insights into microbial community dynamics. However, these methods struggle with the inherent missingness and asynchronous sampling commonly present in longitudinal microbiome data, limiting their predictive accuracy and generalizability in real-world applications.^10-12^ While some studies have attempted to address these issues with statistical spline estimation and dynamic time warping (DTW),^12^ these approaches rely on data interpolation, which could introduce additional biases and distortions. Furthermore, most frameworks remain narrowly tailored to specific ecosystems, restricting their translational potential across diverse clinical and environmental settings.

Meanwhile, advances in sequence modeling, particularly Transformer architectures featuring self-attention and temporal positional encoding, have revolutionized forecasting in fields such as gene expression and single-cell dynamics.^13^ These approaches offer new avenues for learning from incomplete, heterogeneous biological data without explicit imputation. Despite their promise, the systematic application of transformer-based models to longitudinal microbiome analysis—and more broadly, to personalized biological forecasting—remains underexplored.

To address these limitations, we propose MicroProphet, the first personalized digital twin framework leveraging a transformer architecture designed specifically for modeling microbial community dynamics from incomplete longitudinal data. Departing from conventional interpolation-based strategies, MicroProphet directly learns from irregularly sampled data, capturing long-range dependencies and critical turning points in microbial trajectories. Remarkably, the model can forecast subject-specific microbial abundance trajectories using only the initial 30% of time points. We systematically evaluated MicroProphet across diverse ecological settings, including the human gut and skin microbiome, synthetic communities, and corpse decomposition, demonstrating its robust generalization across environments and individuals. Beyond microbial applications, our framework establishes a generalizable approach for real-time monitoring of biological systems, early disease detection, and precision-guided therapeutic interventions, thereby advancing the frontiers of personalized medicine and ecological management.

## Results

### Overview of MicroProphet

MicroProphet is able to generate reliable forecasts of microbial community dynamics from longitudinal microbiome data, even with irregular sampling intervals (see **Methods, Figure 1a, b**). The process begins with slicing the input microbiome time-series data (**Figure 1b-1**), followed by instance normalization and patching to ensure data stability and consistency (**Figure 1b-2**). The data is then mapped into a high-dimensional space, where temporal positional encoding embeds temporal information at each time point, allowing MicroProphet to recognize temporal context (**Figure 1b-3**). The transformer encoder utilizes the multi-head self-attention mechanism to capture long-term dependencies in the time series and extract dynamic features from the microbial community (**Figure 1b-4**). Finally, the model flattens the data and passes it through a linear layer (**Figure 1b-5**) to generate predictions, outputting the predicted microbial abundance for each time point (**Figure 1b-6**). By modeling temporal relationships, MicroProphet provides detailed insights into microbial fluctuations, enabling personalized predictions of microbial dynamics for individual subjects **(Figure 1c**) and anticipating subject-specific health outcomes (**Figure 1d**). Beyond prediction, the framework aids in identifying disease-associated microbial signatures (**Figure 1e**). The attention mechanism captures critical temporal dependencies and identifies key transition timepoints in microbial dynamics (**Figure 1f**), enabling real-time monitoring of microbial community shifts. Additionally, MicroProphet supports the reconstruction of ecological interaction networks (**Figure 1g**), offering a systems-level understanding of microbial dynamics across diverse environments.

**Figure 1.**
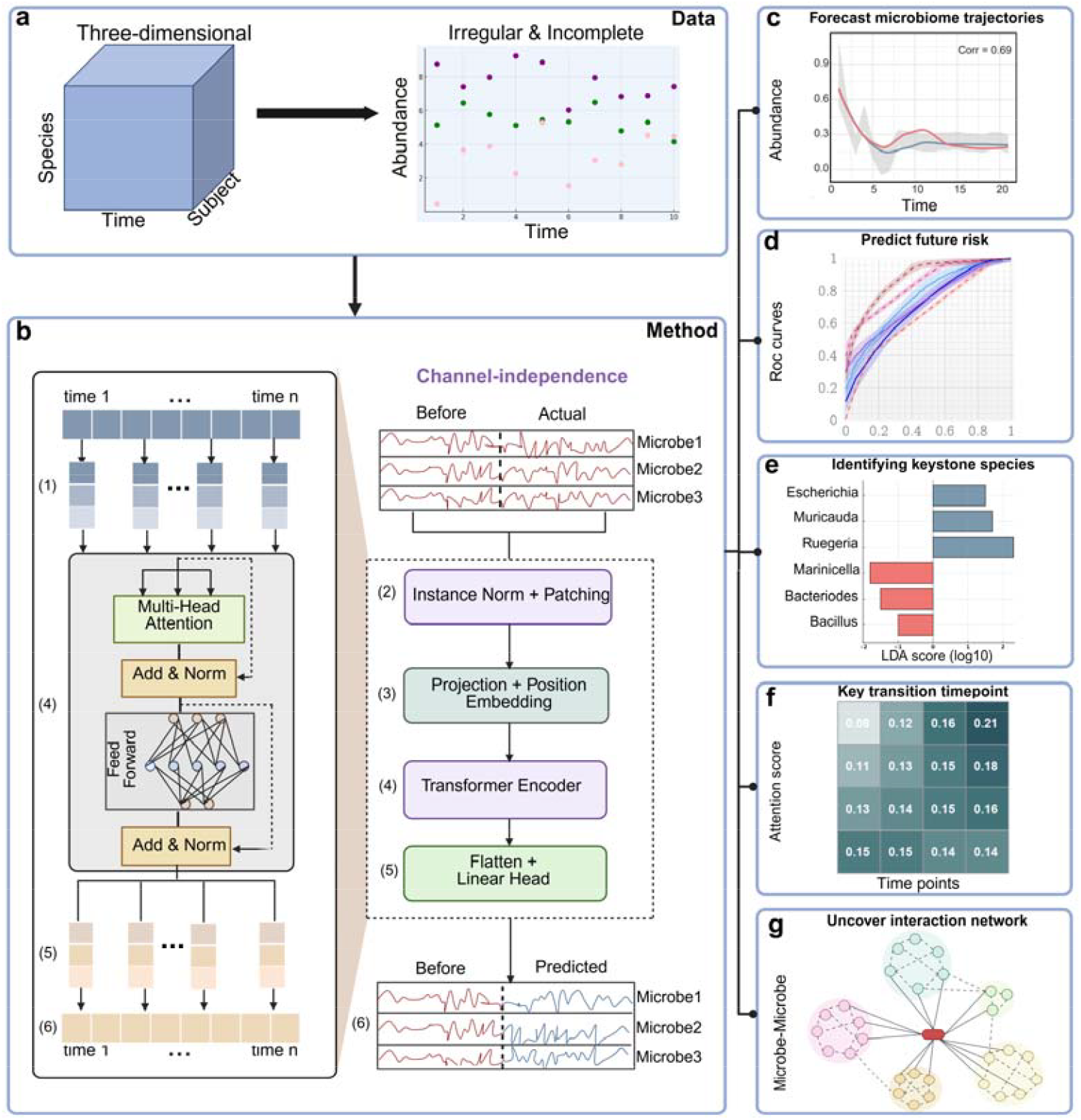
Overview of MicroProphet framework. (a) The structure of the longitudinal microbiome data. The data are represented as a three-dimensional matrix consisting of species, time, and subject dimensions. Due to inconsistent sampling time points across subjects, the data exhibit a missing and irregular distribution. **(b) The overall architecture of MicroProphet framework**. The input consists of multiple univariate time series representing microbial abundances. Instance normalization and patching (Instance Norm + Patching) are applied for data preprocessing. The core of the framework is a Transformer encoder, which captures dynamic relationships between microbes. The encoder includes a multi-head attention mechanism and a position embedding module. The output is generated via a linear head after flattening. **(c-g) Applications of MicroProphet in different contexts**. c, real-time monitoring of microbial dynamic trajectories; d, predicting an individual’s future health status; e, identifying keystone microbial species; f, identifying key transition timepoints using attention mechanism; g, inferring interaction networks among microbes.

**Figure 2.**
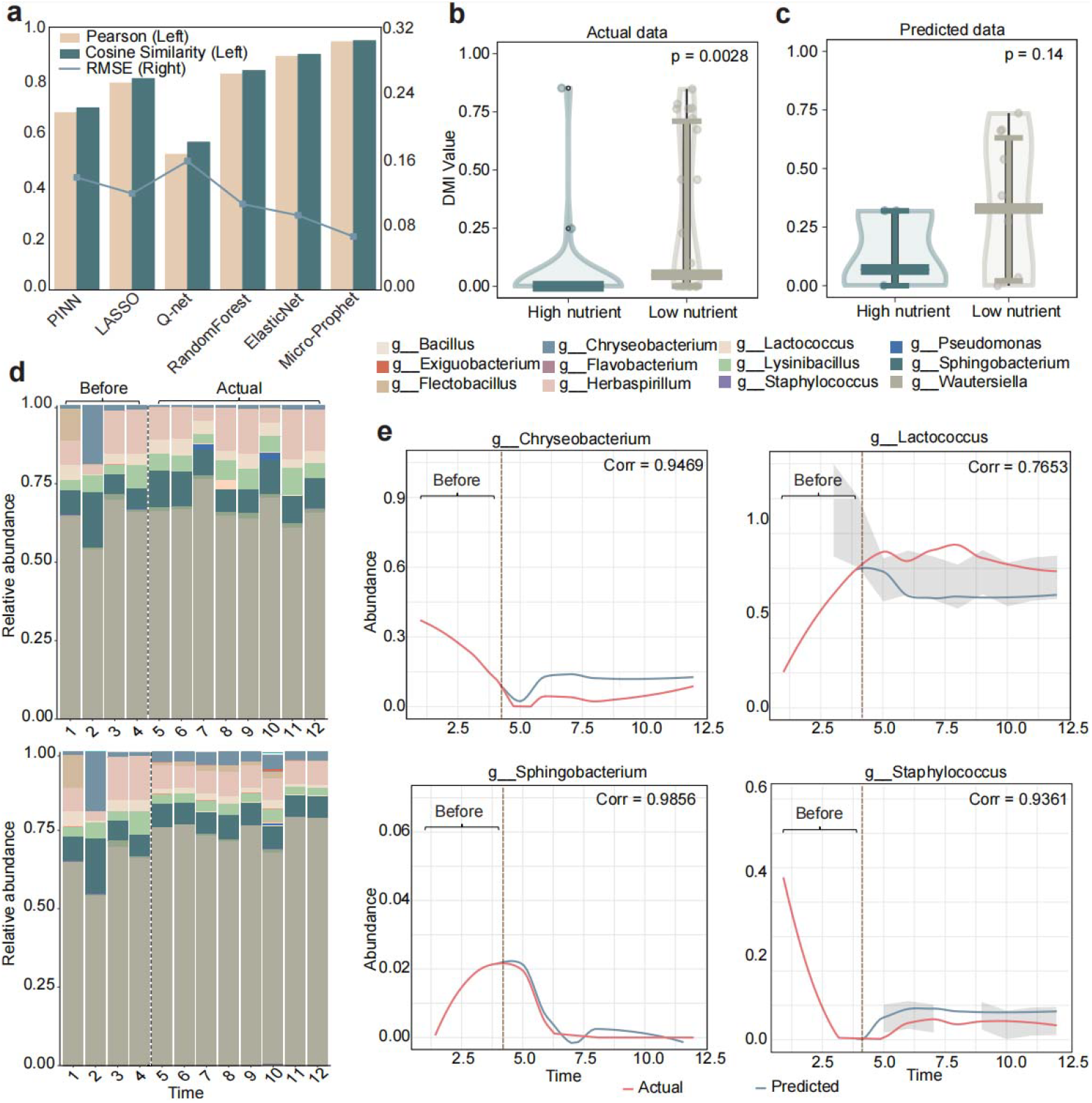
Performance evaluation of MicroProphet on the Synthetic Microbial Communities dataset. **(a) Comparison of predictive performance among different models**. Bar plots represent Pearson correlation (yellow) and cosine similarity (green), while the line plot represents RMSE (blue). **(b-c) Distribution of the DMI value in high-nutrient and low-nutrient communities**. b, DMI for actual data, with a significant difference (p = 0.0028); c, DMI for predicted data, with a p-value of 0.14. Blue represents high nutrient, and green represents low nutrient. **(d) Stacked bar charts showing microbial composition over time**. The top panel represents the actual observed microbial data, while the bottom panel represents the predicted data. The horizontal axis represents time, and the vertical axis represents microbial abundance. **(e) Predicted trajectories of highly abundant microbial taxa**. Line plots show the temporal abundance trajectories of four representative microbial taxa for a single subject. The red lines represent the actual abundances, while the blue lines represent the predicted abundances. Shaded areas indicate confidence intervals for the predictions. The x-axis represents time, and the y-axis represents microbial abundance values. RMSE, root mean squared error; DMI, degree of microbial individuality.

**Figure 3.**
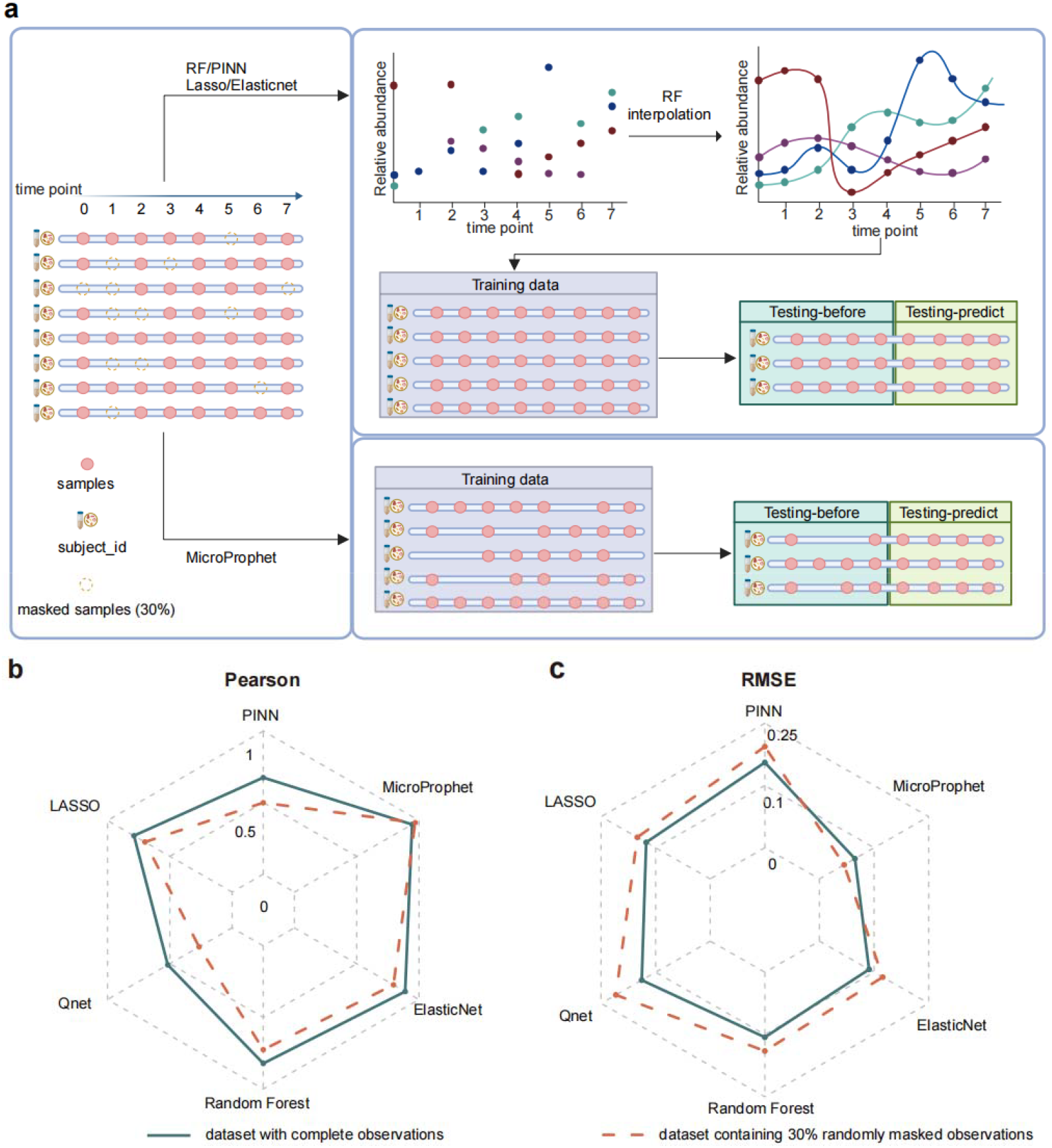
Evaluating model robustness under randomly masked observations. **(a) Schematic illustrating the 30% random masking strategy applied to the Synthetic Microbial Communities dataset**. The top panel illustrates the workflow for benchmark methods (e.g., PINN, LASSO, ElasticNet, Q-net, and Random Forest), which require performing RF-based interpolation to fill in missing observations caused by the 30% random masking before proceeding with prediction. While the bottom sections depict how training and testing are conducted for MicroProphet, which directly handles incomplete dataset without interpolation. **(b-c) Comparison of predictive performance between MicroProphet and benchmark methods on the dataset with complete observations (green solid line) and dataset containing 30% randomly masked observations (red dashed line)**. (b) Pearson correlation coefficient (higher values indicate better prediction accuracy); (c) Root mean squared error (RMSE; lower values indicate better prediction accuracy).

**Figure 4.**
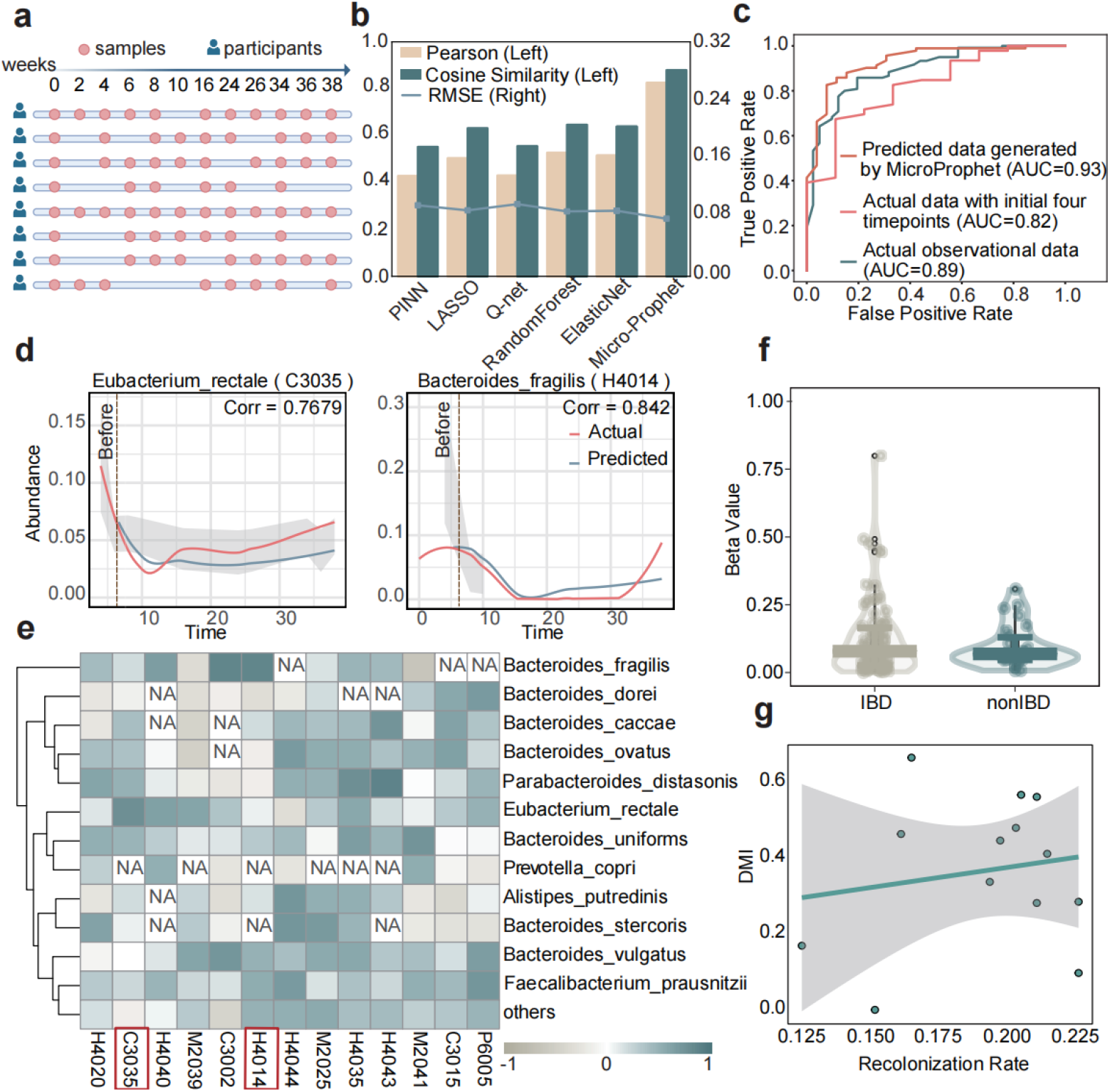
Microbial abundance predictions using the MicroProphet model on the IBD dataset. **(a) Sampling time points for several patients in the IBD dataset**. Each row represents a patient, and red dots indicate the actual measured time points. **(b) Comparison of prediction performance for different methods**. The left panel shows Pearson and Cosine Similarity metrics (in yellow and green respectively), while the right panel shows RMSE (in blue). **(c) Classification performance for distinguishing IBD from non-IBD subjects**. Employed random forest models to distinguish between IBD patients using three distinct datasets: the data predicted by MicroProphet combined with actual samples from the initial four time points (orange); actual samples only from the initial four time points (red); the actual observational dataset (green). **(d) Predicted trajectories of *Bacteroides uniformis* and *Eubacterium rectale***. Line plots show the temporal abundance trajectories of *Bacteroides uniformis* and *Eubacterium rectale* for a single subject. The red lines represent the actual abundances, while the blue lines represent the predicted abundances. Shaded areas indicate confidence intervals for the predictions. The x-axis represents time, and the y-axis represents microbial abundance values. **(e) Pearson correlation heatmap between actual and predicted microbial abundances for top-ranked species across subjects**. Darker colors indicate higher correlations, whereas lighter colors represent lower correlations. “NA” values indicate cases where the microbial abundance was zero, making Pearson correlation calculation unavailable. **(f) Linear regression analysis of microbiome stability over time**. Blue represents non-IBD subjects, and green represents IBD subjects. Y-axis representing the Beta values from the analysis. **(g) Correlation between the DMI index and bacterial recolonization rate**. The x-axis represents the value of recolonization rate, and the y-axis represents the DMI value. RMSE, root mean squared error; DMI, degree of microbial individuality.

**Figure 5.**
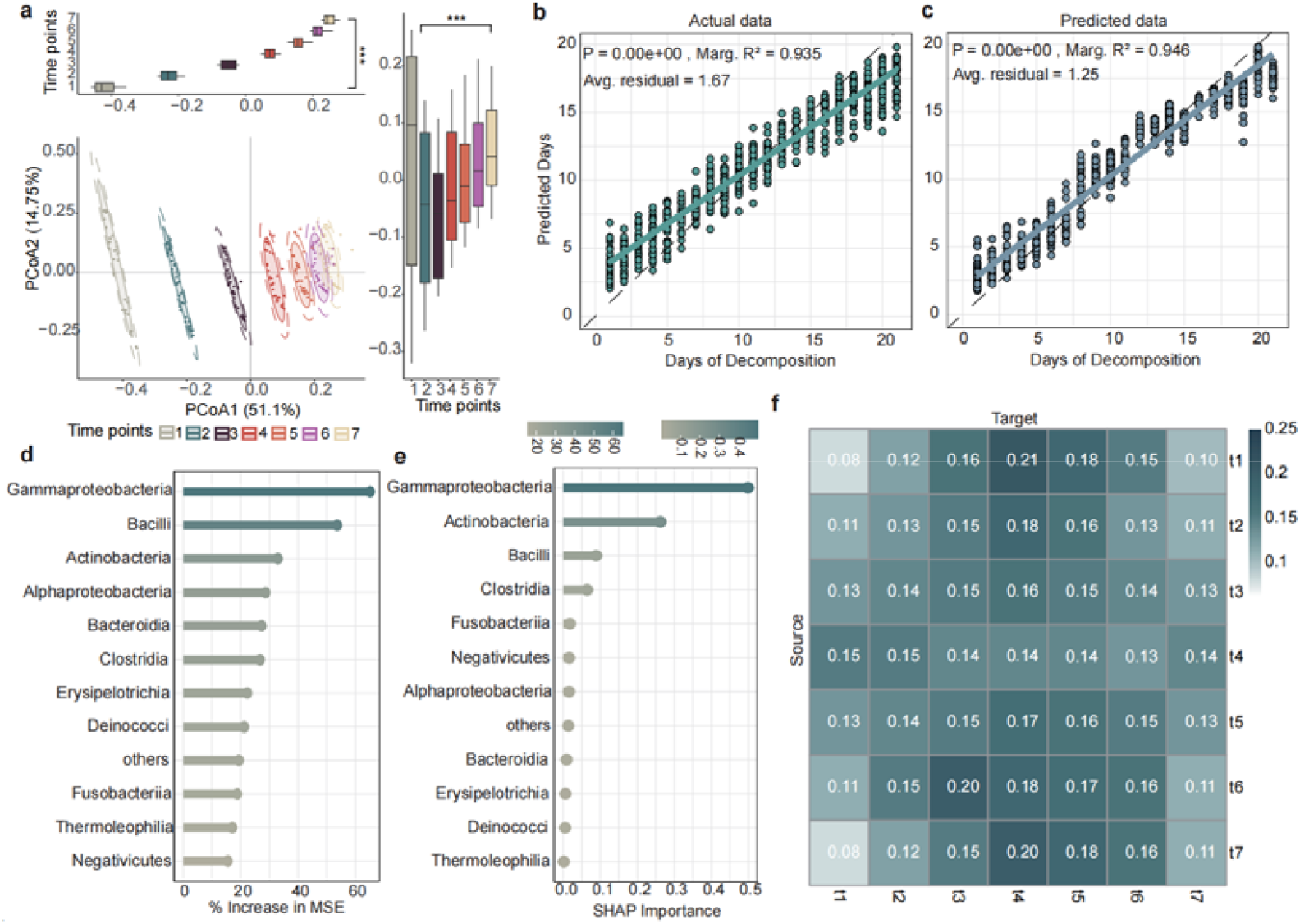
Microbial abundance predictions using the MicroProphet model on the Corpse dataset with complete time points. **(a) Beta diversity analysis at initial time points**. The clear separation between time points along the PCoA1 axis indicates strong temporal shifts in microbial community composition during decomposition. Each point represents a microbial sample, and colored ellipses indicate 95% confidence intervals for each time point. Group differences were tested using permutational multivariate analysis of variance (PERMANOVA), with *** indicating p < 0.001. **(b) Linear regression of decomposition days using actual microbial data**. The scatter plot illustrates the regression of actual days of decomposition based on the microbial abundance data using a Random Forest model. Y-axis represents predicted days; X-axis represents decomposition days. **(c) Linear regression of decomposition days using predicted microbial data**. The scatter plot demonstrates the regression of decomposition days using the predicted microbial abundance data from the MicroProphet model. The Y-axis represents predicted days; the X-axis represents decomposition days. **(d) Feature importance ranking of microbial taxa based on the random forest model for days of decomposition prediction**. The horizontal axis represents the percentage increase in mean squared error (% increase in MSE). **(e) Feature importance ranking derived from SHAP analysis in the MicroProphet model**. The horizontal axis represents the standardized SHAP values. **(f) Attention scores among different time points**. Each cell represents the attention score computed by the MicroProphet from one initial time point (rows) to another (columns). The color intensity indicates the strength of the attention, with blue representing stronger attention and yellow representing weaker attention. For example, t1 to t4 shows the strongest attention score (0.21), indicating that t1 has the highest dependency on t4 during the prediction process.

**Figure 6.**
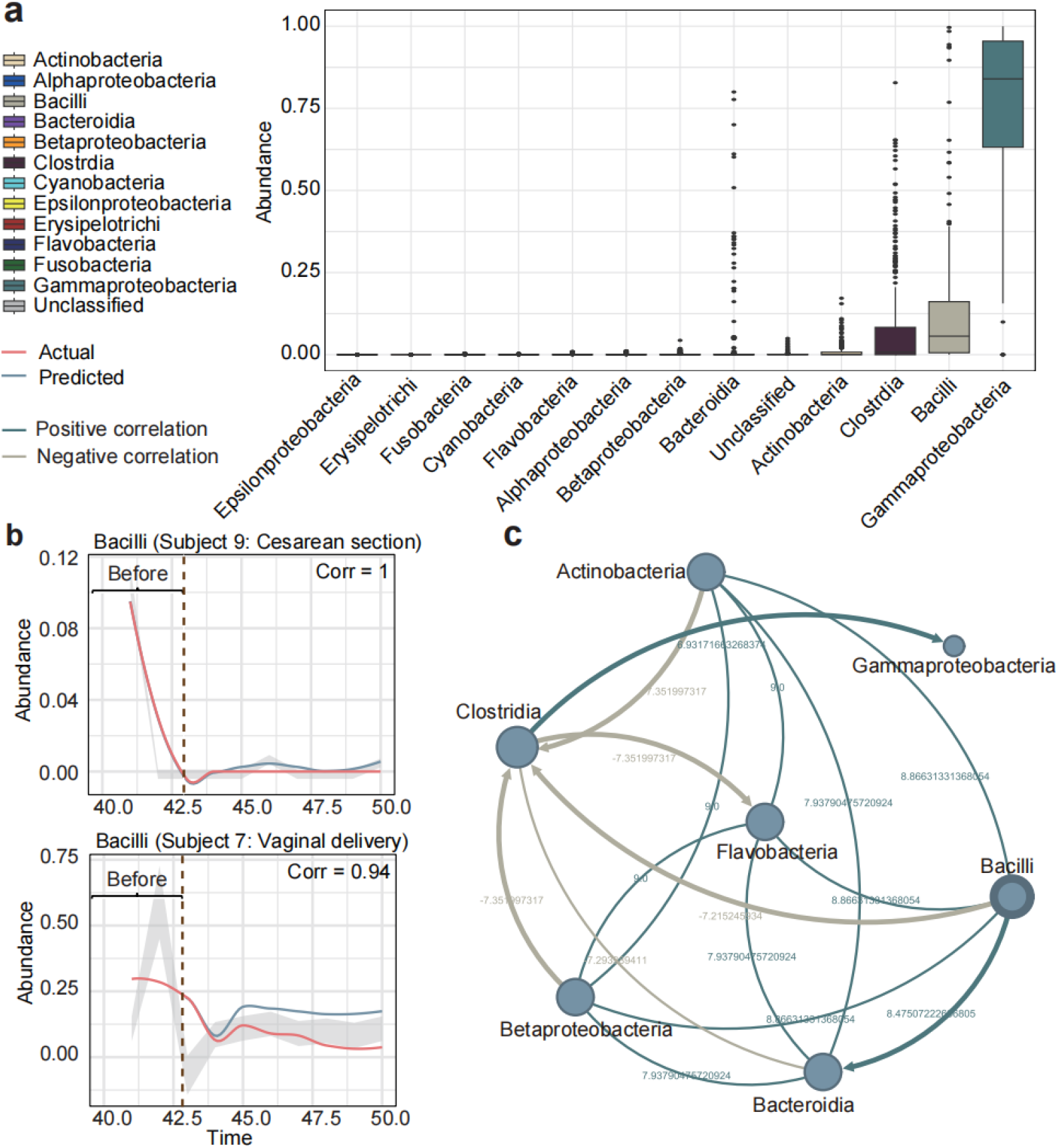
Personalized trajectories and microbial interactions in infant gut microbiome. (**a) The abundance distribution of different microbial taxa in the infant gut microbiome**. (**b) Predicted microbial abundance trajectories for Bacilli in infants born via cesarean section and vaginal delivery**. The red line represents actual microbial abundance, while the blue line represents predicted values. Pearson correlation coefficients are displayed for each trajectory. The x-axis represents time, and the y-axis represents microbial abundance values. (**c) Microbial interaction network based on LSA**. Light blue lines represent positive correlations, while light brown lines represent negative correlations between the bacterial groups. The values on the edges represent the correlation coefficients. he direction of the relationship is determined by the Kolmogorov-Smirnov test, with a p-value < 0.05 indicating the presence of a directed relationship. LSA, Local Similarity Analysis.

To evaluate the performance of MicroProphet, we applied it to four publicly available longitudinal microbiome datasets representing diverse microbial ecosystems (**Supplementary table 1**): (1) the Synthetic Microbial Communities dataset,^14^ consisting of 384 samples from 32 artificial communities constructed from subsets of 50 microbial isolates over a 12-day period; (2) the inflammatory bowel disease (IBD) dataset,^15^ from the Integrative Human Microbiome Project (iHMP), containing 534 longitudinal stool samples from 65 individuals (IBD patients and healthy controls); (3) the Corpse dataset,^16^ comprising 21 time-point samples collected from the face skin of 34 decomposing human cadavers at three U.S. forensic research sites; (4) the Infant dataset,^17^ including 360 aligned metagenomic samples from 36 preterm infants collected across 10 time points. Together, these datasets enable rigorous assessment of MicroProphet across both human-associated and environmental microbiome contexts, with varying degrees of temporal

### Predicting microbial abundance trajectories across the whole time series in synthetic microbial communities

To systematically evaluate the performance of MicroProphet, we applied it to the Synthetic Microbial Communities dataset with complete and regularly spaced time points.^18^ This dataset comprises 32 synthetic microbial communities, each sampled at 12 distinct time points with full observed measurements (see **Methods, Supplementary table 1**). We first assessed MicroProphet’s prediction accuracy by varying the fraction of initial time points from 10% to 50%. Notably, the prediction performance improved as the proportion of known data increased, and the accuracy began to saturate around the initial 30% of time points (**Supplementary Figure 2a, Supplementary table 2**). This finding demonstrates MicroProphet’s ability to achieve reliable predictions even with limited data.

Next, we compared the performance of MicroProphet with five benchmark methods, including PINN (a Physics-Informed Neural Network employing LSTM to solve the generalized Lotka-Volterra equations^19^), Q-net^20^ (a digital twin model based on conditional inference trees), LASSO, ElasticNet, and Random Forest (**Figure 2a**).

Specifically, the data were divided into a training set (70%) and a test set (30%). We then used the initial 30% of time points to predict the subsequent 70%. Overall, MicroProphet demonstrated superior performance, achieving a Pearson correlation coefficient of 0.9447, a cosine similarity of 0.9487, and a root mean squared error (RMSE) of 0.065 (**Supplementary table 3**). Taking one microbial community (ID: 7F) as an example, the predicted microbial abundance distribution closely matched the observed distribution (**Figure 2d**), indicating that the model accurately captured the dynamic changes of individual genus over time (**Figure 2e**, Pearson > 0.75).

Beyond abundance prediction, we assessed microbial community stability at the genus level under high-nutrient and low-nutrient conditions (see **Methods**), hypothesizing that stability might be environment-specific. To quantify this, we introduced the degree of microbial individuality (DMI), a metric that measures the similarity within an individual relative to the population (see **Methods**). A higher DMI indicates greater microbial individuality. Our results indicated that the DMI value for microbes was significantly lower in high-nutrient communities (Mann-Whitney U test, p = 0.0028, **Figure 2b, Supplementary table 4**), supporting the hypothesis that high-nutrient environments are more prone to microbial invasion.^18^ Notably, the same trend was observed in the predicted dataset (**Figure 2c**), further validating the robustness of MicroProphet in capturing microbial community dynamics from both an ecological and theoretical perspective. This comprehensive evaluation underscores that MicroProphet not only predicts microbial abundance accurately but also provides insights into community stability and dynamics under different conditions.

### Evaluating the robustness of MicroProphet in handling missing observations through random masking

To assess MicroProphet’s robustness in handling incomplete datasets that contain missing observations, we simulated an irregular sampling scenario by randomly masking 30% of the samples in the Synthetic Microbial Communities dataset (**Figure 3a, Supplementary table 1**). Traditional methods, such as Lasso, ElasticNet, and Random Forest, typically address missing observations through data interpolation (e.g., Random Forest interpolation) prior to predictions. However, we observed that interpolation introduced significant discrepancies between imputed and true values, potentially biasing subsequent predictions (**Supplementary Figure 3**). In contrast, MicroProphet directly modeled incomplete datasets without interpolation, consistently maintaining high predictive accuracy (Pearson correlation = 0.969, cosine similarity = 0.970, RMSE = 0.05, **Supplementary table 3**). On the other hand, the performance of benchmark methods significantly declined when applied to incomplete datasets (**Figure 3b-c**), reinforcing the effectiveness of MicroProphet in preserving prediction accuracy.

Further evaluations confirmed that MicroProphet’s performance remained robust even as the proportion of missing observations increased. When progressively increasing the proportion of missing observations from 0% to 60%, prediction accuracy gradually declined, yet MicroProphet still maintained strong performance at a missing data proportion of 30% (**Supplementary Figure 2b, Supplementary table 2**). These findings underscore MicroProphet’s exceptional capability to handle incomplete longitudinal microbiome data, positioning it as a highly reliable tool for forecasting microbial community dynamics in realistic scenarios characterized by irregular sampling.

### MicroProphet enables personalized prediction of microbial community dynamics in the IBD cohort

Next, we validated MicroProphet’s ability to generate personalized microbial trajectory forecasts using the IBD dataset^15^ (**Supplementary table 1**). The dataset comprises longitudinal microbiome profiles from 65 participants, including samples collected over 38 weeks, with subjects grouped into IBD and non-IBD categories. Due to its incompleteness nature, this dataset provided an ideal scenario to evaluate both the overall predictive accuracy and MicroProphet’s ability to capture detailed, individualized temporal patterns (**Figure 4a**).

We first compared the predictive performance of MicroProphet against other methods. MicroProphet consistently demonstrated superior performance across all evaluated metrics (**Figure 4b, Supplementary table 3**). Specifically, we calculated the Pearson correlation between the actual and predicted abundances for each top-ranked microbial species across all test subjects. Most individuals achieved highly accurate predictions (**Figure 4e**). For instance, *Eubacterium rectale*, a species with anti-inflammatory properties often depleted in IBD patients, and *Bacteroides fragilis*, which plays a dual role in gut health and IBD pathogenesis, exhibited distinct temporal variation patterns in both actual and predicted data (**Figure 4d**). ^21^ Moreover, beta-diversity analysis indicated no significant differences between predicted and actual microbial compositions (**Supplementary Figure 4**). To further support the model’s clinical utility, we employed random forest models to distinguish IBD patients using three distinct datasets: (1) the data predicted by MicroProphet combined with actual samples from the initial four time points, (2) actual samples only from the initial four time points, and (3) the actual observational dataset (**Figure 4c**). Remarkably, the random forest model trained on data predicted by MicroProphet achieved classification accuracy comparable to that the actual observational dataset, significantly outperforming predictions based solely on data from initial four time points. These results underscore MicroProphet’s ability to accurately infer future health status from limited temporal microbiome data.

To further simulate the dynamic changes in microbial abundance trajectories under different intervention effects, we selected one IBD patient who had not taken antibiotics (C3037) and another who had taken antibiotics (P6016) from the training dataset to fit their respective trajectory changes (**Supplementary Figure 5**). We observed that although both subjects were diagnosed with IBD, the two key species (*Bacteroides uniformis* and *Eubacterium rectale*) exhibited markedly different trajectories due to antibiotic intervention. Specifically, patient P6016 showed a pronounced decline in microbial abundance at early stages, reflecting the antibiotic treatment effect. These personalized predictions not only reveal subtle temporal variations that may indicate disease progression or therapeutic response, but also highlight the potential of MicroProphet for proactive and intervention strategies.

Additionally, the longitudinal data allow us to investigate microbial community stability by quantifying the dissimilarity between-sample pairs in relation to collection date intervals. Our analysis revealed that the IBD-associated gut microbiome exhibited a more higher rate of temporal fluctuation compared to non-IBD microbiomes (**Figure 4f**), corroborating previous reports.^15^ Such microbiome instability may play a critical role in IBD onset, progression, and inflammatory regulation, further suggesting the microbiome as a viable target for personalized interventions. Meanwhile, we assessed the species recolonization rate, measured by consistency when a species is detectable after being undetected in one 1 − pairwise Jaccard distance)^22^ was significantly associated with DMI or more consecutive samples. The overall recolonization rate (measured by (**Supplementary table 3 and 5**). Our results suggested that highly individualized strains are more likely to recolonize (**Figure 4g**), validating a hypothesis raised by fecal transplantation studies^8^ and implying its potential efficacy in treating IBD-associated dysbiosis.

Collectively, these findings demonstrate that our transformer-based digital twin framework not only accurately predicts longitudinal microbial dynamics in the IBD cohort but also offers promising applications for personalized microbiome-based healthcare. By enabling real-time monitoring of microbial responses to therapeutic interventions, MicroProphet represents a powerful computational tool to support predictive intervention strategies in managing chronic diseases like IBD.

### Uncovering microbial dynamics during corpse decomposition with MicroProphet

To demonstrate MicroProphet’s capability to establish reliable long-term forecasts from limited initial observations, we applied it to the Corpse dataset (**Supplementary table 1**), modeling microbial dynamics during human corpse decomposition and fitting decomposition timelines based on predicted data. The dataset ^23^ was obtained from the face skin of 34 corpses sampled continuously over 21 days of decomposition.

Beta diversity analysis at initial 30% of time points (**Figure 5a**) revealed clear temporal shifts in microbial community composition, with a marked separation along the PCoA1 axis. This pattern indicates substantial microbial changes during early decomposition, followed by a progressive stabilization after day 4 of decomposition. MicroProphet achieved high accuracy in predicting microbial abundance trajectories, demonstrating strong performance across multiple evaluation metrics (Pearson correlation = 0.72; cosine similarity = 0.776; RMSE = 0.118; **Supplementary Figure 6 and Supplementary table 3**). The predicted temporal trajectories closely mirrored the actual observed microbial dynamics, accurately capturing the characteristic rise and fall of microbial taxa abundances during decomposition (**Supplementary Figure 7**). To further validate these predictions, we employed random forest regression models trained separately on actual and predicted microbial abundance data to estimate decomposition days. Both models achieved high regression accuracy (R^2^ > 0.9; **Figure 5b, c**), underscoring MicroProphet’s reliability in accurately forecasting decomposition progression and estimating days of decomposition.

Next, we compared the feature importance rankings derived from the random forest model for estimating days of decomposition (**Figure 5d**) with the microbial feature importance calculated by SHAP analysis in MicroProphet (**Figure 5e**, see **Methods**). The results showed a high degree of consistency between the two methods in identifying key microbial taxa. Notably, Gammaproteobacteria, Bacilli, and Actinobacteria ranked among the top taxa in both approaches, supporting previous findings on their involvement in corpse decomposition.^24,25^ Gammaproteobacteria exhibited a significant increase during the early and middle stages of decomposition, followed by a decline in the late stage (**Supplementary Figure 7**). This initial proliferation is likely associated with its ability to thrive in low-oxygen or anaerobic conditions while secreting hydrolytic enzymes such as proteases and lipases, which facilitate the breakdown of soft tissues^26^. However, as decomposition progresses, the depletion of tissue substrates and shifts in environmental conditions may limit its further growth, leading to a reduction in abundance. This decline may signal the transition from tissue degradation to the later mineralization stage of decomposition. Bacilli, widely present in human skin, soil, and external environments, possess a sporulation ability that confers resistance to extreme conditions, allowing them to persist even in the late stages of decomposition^26^ (**Supplementary Figure 7**).

To investigate the temporal dynamics and dependencies between time points, we analyzed the attention scores generated by MicroProphet. The results revealed that, when predicting future microbial abundances trajectories, initial time points exhibited a notably higher dependency on the fourth time point. Specifically, its cumulative attention score (1.22) was as high as 54.4% above that of time point 7, and as low as 8.9% above that of time point 5 (**Figure 5f**). We thus infer that day 4 of decomposition may represent a critical transition point in the microbial succession process. This finding, together with the observed stabilization in beta diversity after day 4 of decomposition (**Figure 5a**), which indicated substantial shifts in community composition during the initial four days, followed by a progressive stabilization thereafter.

Ultimately, the model’s ability to capture long-term microbial trends and accurately predict decomposition days underscores its potential as a powerful tool for forensic applications. Notably, the attention mechanism identified crucial transitional time points, such as day 4 of decomposition, which may serve as inflection points in microbial community succession.

### Personalized predictions of infant gut microbial dynamics across cesarean and vaginal births

MicroProphet could also be applied to modeling dynamic changes in the early infant gut microbiome, which are characterized by high irregularity and pose significant challenges for accurate prediction. To capture the highly unstable individual-level microbial shifts, we simulated personalized microbial abundance trajectories for 36 infants using microbiome data collected over a 40 to 50-day period after birth (**Supplementary table 1**). Initially, infant gut microbial communities were dominated by Clostridia, Bacilli, and Gammaproteobacteria (**Figure 6a**), which formed the core microbial community. Using MicroProphet, we predicted the microbial abundance trajectories for each infant individually, capturing the temporal dynamics of the gut microbiome in a personalized manner. As shown in **Figure 6b**, the predicted trajectories of Bacilli differed between infants born via cesarean section (C-section) and vaginal delivery (VD). Specifically, infants delivered vaginally exhibited a consistently higher abundance of Bacilli compared to infants delivered via C-section (**Figure 6b**), aligning with previous reports suggesting that vaginal birth exposes infants directly to maternal microbiota, influencing early microbial colonization patterns ^27^.

To further elucidate the temporal interactions within the infant gut microbiome, we employed Local Similarity Analysis (LSA),^28^ a method capable of capturing time-lagged correlations that may be overlooked by Pearson correlation analysis (**Supplementary table 6**). We identified Bacilli and Clostridia as key nodes within the microbial interaction network. According to previous studies,^29,30^ Bacilli, commonly associated with early gut colonization, play a crucial role in nutrient metabolism and immune system modulation. Clostridia, on the other hand, are prominent members of the gut microbiota and are known for their role in fermenting complex carbohydrates, producing short-chain fatty acids, and maintaining gut homeostasis.^31^ We employed Kolmogorov-Smirnov (KS) testing to determine the directional relationships between microbes, assessing whether one microbe statistically preceded another in terms of its distribution. For example, we observed a directed positive correlation where the temporal distribution of Clostridia preceded that of Flavobacteria (**Figure 6c**). Such temporal precedence suggested Clostridia might create environmental conditions favorable for subsequent colonization by Flavobacteria, potentially via metabolic activities such as short-chain fatty acid production.^32^ These nuanced temporal microbial interactions highlight MicroProphet’s sensitivity and accuracy in capturing biologically meaningful, dynamic microbiome relationships.

Taken together, these results illustrate MicroProphet’s significant potential for personalized forecasting of early-life gut microbiome dynamics, providing novel insights into microbial colonization patterns influenced by birth mode. This capability holds promise for guiding precise, personalized microbiome-targeted interventions during the critical early developmental window, thereby promoting long-term infant health.

## Discussion

Despite growing recognition of the importance of forecasting microbial community dynamics in both clinical and ecological contexts, current modeling approaches are hampered by persistent limitations in handling real-world longitudinal data. Traditional methods often depend on interpolation or data alignment to handle missing observations, procedures that may distort true microbial temporal signals and reduce biological fidelity. Moreover, these conventional methods, such as the gLV equations, rely on rigid assumptions about microbial interactions, restricting their applicability across diverse microbiome scenarios. Consequently, such methodological constraints hinder the development of accurate and actionable forecasts, which are essential for timely clinical interventions or ecological management. Here, we address these challenges by proposing MicroProphet, a generative digital twin framework that captures complex temporal dependencies and exhibits adaptability across multiple scenarios.

One of the key findings from this study is the ability of MicroProphet to make personalized predictions of microbial dynamics, particularly in the clinical contexts like IBD. In contrast to traditional methods that assume one-size-fits-all approaches to microbial data, our framework can uncover subtle microbial shifts uniquely tied to each patient’s health status and disease progression. For instance, *Faecalibacterium prausnitzii*, an anti-inflammatory bacterium, is typically reduced in abundance in IBD patients. ^33,34^ Conversely, *Bacteroides fragilis*, associated with pro-inflammatory responses, is often found at higher levels in these patients.^35^ By accurately forecasting the abundance of such key taxa, MicroProphet can predict disease states like IBD with high accuracy (AUC = 0.93), highlighting its utility in identifying early, personalized biomarkers and disease markers. This discovery lays the foundation for future intervention studies. MicroProphet’s predictive capacity could guide the timing of fecal microbiota transplantation (FMT) or probiotic administration (e.g., Lactobacillus and Bifidobacterium-based formulations) based on real-time microbial trajectory simulations.^36,37^ The subject-specific forecasts provided by MicroProphet could inform the optimal timing and dosage of such interventions, thereby moving us closer to truly personalized, microbiome-based therapies that are both accurate and clinically relevant.

Another critical aspect of MicroProphet is its capacity to handle incomplete dataset. Most traditional models, including DBN^12^ and MDSINE^7^, require data alignment or interpolation, processes that can introduce bias or remove valuable information, especially when dealing with missing observations. MicroProphet, however, directly processes incomplete longitudinal data without the need for such preprocessing by employing time-aware positional encoding. This feature is handy in real-world settings, where sampling schedules are often irregular. It not only enhances the robustness of predictions but also offers a more accurate representation of microbial community evolution across diverse ecosystems. This flexibility allows MicroProphet to be applied broadly across multiple environments—from human microbiomes to environmental and synthetic microbial communities—without the limitations imposed by rigid data interpolation requirements.

Furthermore, MicroProphet demonstrated robust long-term forecasting capabilities by utilizing only the initial 30% of time points. This reliable predictive performance is enabled by its attention mechanisms, which also effectively capture critical time points indicative of significant microbial community shifts. For example, in the corpse decomposition context, our model successfully predicted microbial community shifts throughout a 21-day decomposition period based solely on microbial abundances measured during the first 7 days. Additionally, the attention mechanism identified crucial transitional time points, such as day 4 of decomposition, which may represent key inflection points in microbial community dynamics. Such predictive accuracy is particularly valuable in forensic microbiology, where estimating the days of decomposition precisely and rapidly is crucial for investigative timelines.^16^ Additionally, early forecasting of microbial succession patterns can substantially benefit environmental microbiology, aiding in the timely detection of ecological disturbances and facilitating proactive interventions. Collectively, these findings underscore the broad potential utility of MicroProphet across various time-sensitive fields by providing accurate, early, and actionable predictions based on minimal initial data. ^3,16^

MicroProphet’s cross-ecosystem adaptability further cements its position as a robust and broadly applicable model for microbial dynamics. In our evaluations, MicroProphet consistently achieved Pearson correlation coefficients exceeding 0.80 and RMSE values below 0.10 across multiple ecosystems, including human gut and skin microbiota, synthetic communities, and corpse decomposition scenarios. Even in the challenging context—predicting community shifts in preterm infants’ gut microbiome—MicroProphet maintained a Pearson correlation of 0.92, underscoring its resilience in highly dynamic and complex environments. By accurately capturing system-specific patterns, the model reveals key ecological factors—such as community stability, recolonization dynamics, and keystone taxa—that inform targeted interventions. This level of predictive power across diverse use cases not only highlights MicroProphet’s robustness but also enables deeper insights into how microbial communities respond to perturbations. Coupled with its capability of handling missing observations, MicroProphet thus provides researchers with the flexibility to identify critical microbial biomarkers and interactions across a wide range of ecological and clinical settings, driving forward precision strategies in both healthcare and environmental management.

Despite these promising advances, our study has certain limitations. While MicroProphet effectively processes microbial abundance data, future work should consider integrating multi-omics datasets (e.g., metagenomics, metabolomics) to provide a more holistic view of host-microbiome interactions. In addition, incorporating real-time monitoring and dynamic feedback mechanisms could further enhance the model’s applicability in clinical decision-making and environmental management. Looking ahead, expanding the model to integrate spatiotemporal data—capturing not only temporal trends but also spatial variations in microbial composition across different body sites or environmental niches—will offer a more comprehensive digital twin of microbial ecosystems, further enriching the predictive capabilities of our framework.

In conclusion, MicroProphet provides a powerful and generative digital twin framework for forecasting microbial community dynamics with personalized precision. The framework consistently demonstrates state-of-the-art predictive performance across diverse ecosystems, highlighting its robustness. Additionally, its time-aware position encoding design directly handles incomplete datasets with substantial missing observations (up to 30%) without relying on interpolation, preserving biological fidelity and avoiding bias introduced. Moreover, the framework reliably establishes long-term forecasts using only the initial 30% of time points. Most importantly, MicroProphet achieves personalized microbial trajectory forecasts under various perturbations, facilitating subject-specific insights critical for precision medicine and tailored interventions. Taken together, these advancements position MicroProphet as an innovative solution to transforming complex microbiome data into actionable predictions, driving forward the fields of microbial ecology, personalized healthcare, and environmental management.

## Materials and Methods

### Data collection and pre-processing

Our study collected four public time-series microbiome datasets for testing our method. These datasets cover microbial communities from different microbial ecosystems, covering both human-associated and environmental microbiomes. **Supplementary table 1** summarizes each longitudinal microbiome data set used in this study.

#### Synthetic Microbial Communities dataset

The synthetic microbial communities dataset was collected by Hu *et al*.^14^ They created 32 different synthetic communities using randomly generated subsets of the library of 50 isolates, each consisting of 12 to 20 genera. Microbial community composition was measured via 16S ribosomal RNA (rRNA) amplicon sequencing. These communities were categorized into high-nutrient and low-nutrient conditions, with the low-nutrient condition consisting of 1 g l™1 of yeast extract, 1 g l™1 of soytone, 10 mM sodium phosphate, and trace elements, while the high-nutrient condition was supplemented with 5 g l™1 of glucose and 4 g l™1 of urea to increase interaction strength by amplifying resource competition and promoting environmental pH fluctuations. Overall, 384 samples from 32 synthetic communities over 12 days were used for further analysis.

#### Inflammatory bowel disease (IBD) dataset

The IBD microbiome dataset was collected by Lloyd-Price *et al*. as part of the Integrative Human Microbiome Project (HMP2 or iHMP).^15^ This large-scale study aimed to characterize the gut microbial ecosystem in patients with IBD, including Crohn’s disease (CD) and ulcerative colitis (UC), through a multi-omics approach. Longitudinal stool samples were collected from 132 participants, including IBD patients and healthy controls. Samples were taken every two weeks for one year. A subset of 534 samples from 65 participants were used for the evaluation of MicroProphet. The microbes were identified at the species level.

#### Corpse dataset

This dataset consists of microbial community data collected from 36 human corpses placed outdoors at three willed-body donation facilities: Colorado Mesa University Forensic Investigation Research Station (FIRS), Sam Houston State University Southeast Texas Applied Forensic Science (STAFS) Facility, and University of Tennessee Anthropology Research Facility (ARF).^16^ The corpses were exposed to varying environmental conditions over the years 2016 and 2017. Data on ambient temperature, humidity, scavenging status, and insect activity were also recorded. Daily samples were collected to analyze microbial dynamics during the decomposition process. We utilized the 16S rRNA gene amplicon dataset of face skin for analysis. A subset of 34 corpses sampled at 21 time points was filtered for forecasting, with a focus on 11 microbial taxa (class level).

#### Infant dataset

The dataset was collected by La Rosa *et al*.^17^ from 58 pre-term infants in a neonatal intensive care unit (NICU). The data include 922 gut metagenomes over the first 12 weeks of life, with microbial samples collected every day or two on average. Temporal alignment was performed to address varying rates of change between individuals by Lugo-Martinez *et al*.^12^ The time scale of each sample was warped to match the scale of a reference sample using a first-degree polynomial transformation function. This alignment ensured that corresponding time points across individuals were comparable, enabling accurate comparisons of microbial abundances and improving the identification of dynamic relationships between microbial taxa (class level). A total of 360 gut metagenomes from 36 infants at 10 time points were filtered for forecasting.

### Model Architecture

We developed a temporal modeling framework named MicroProphet for microbial time-series prediction, based on a modified PatchTST (Time Series Transformer) model^38^ implemented using the Huggingface Transformers library. This model comprises two transformer layers, each integrating multi-head self-attention and feed-forward neural networks with Gaussian Error Linear Unit (GELU) activations and layer normalization. Key parameters include the number of input channels (i.e., the number of features in the dataset), context length (i.e., total number of time points minus the forecast horizon), forecast horizon (30% of the total number of time points), number of hidden layers (3 layers), embedding dimension (64), patch length (1 time step), patch stride (1 time step), number of attention heads (4), feed-forward network dimension (256), use of scaling (not applied), and prediction length (equal to the forecast horizon). Training is performed with a learning rate of 10^-4^, batch size of 8, 1000 epochs, early stopping with a patience of 3, and evaluation at the end of each epoch.

#### Data Representation and Patching

Each microbial time series *x*_i_(*t*) is represented as a sequence of abundance values over time. For a given microorganism *i*, the time series *x*_i_ = (*x*_*i*_(1),*x*_*i*_ (2),…,*x*_*i*_(*T*)), where *T* is the total number of observed time points, forms the input for MicroProphet. The data are then divided into patches, where each patch corresponds to a fixed-length window of the time series, denoted by *P*, with a stride *s*. These patches are designed to capture the temporal dependencies of microbial abundance across consecutive time intervals.

Specifically, the patching process generates *M* patches, where:

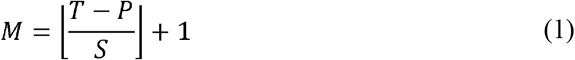

Each patch 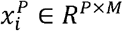 represents a segment of the time series and is treated as an input token to the Transformer model. By applying this patching method, MicroProphet reduces the complexity of long time series data while preserving temporal dependencies, thereby enabling more efficient learning of microbial abundance dynamics. To ensure consistency in input dimensions, padding is applied by repeating the last observed value *x*_*i*_(*T*) at the end of the sequence before patching:

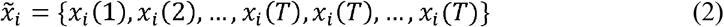

Where the padding length is set to *s*. This approach prevents the model from losing information at the sequence boundary and stabilizes learning.

#### Transformer Encoder

The core of the MicroProphet is the Transformer encoder, which maps the observed microbial abundance sequence into latent representations.

Each patch 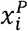 is projected into a high-dimensional space via a linear projection matrix:

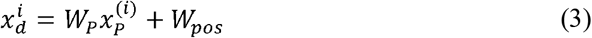

where *W*_*p*_ ∈ *R* ^*D*×*P*^, followed by the addition of positional encoding *R*_*pos*_ ∈ *R*^*P*×*M*^, which encodes the temporal information of each patch.

The transformed patches are then passed through multiple layers of multi-head attention and feed-forward networks. For each head *h*, the attention mechanism computes query, key, and value representations as follows:

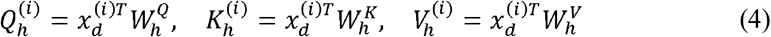

Where 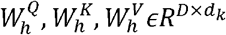 are trainable weight matrices. The scaled dot-product attention is computed as:

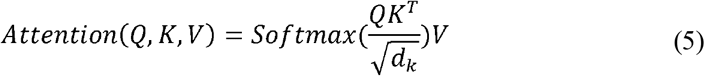

The Transformer model outputs a sequence of latent representations *Z*_*i*_(*t*), which are then used to predict the future abundance of each microorganism 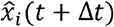, where Δ*t* represents the forecast horizon. The prediction process is performed by applying a linear layer to the output of the Transformer encoder:

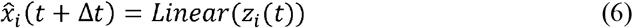

#### Model Training and Loss Function

MicroProphet is trained using a mean squared error (MSE) loss function, which minimizes the difference between microbial abundances 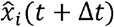 and the ground truth *x*_*i*_(*t* + Δ*t*). The loss is the predicted computed as:

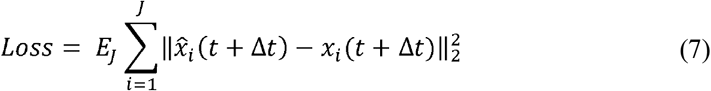

Where *J* is the number of subjects in the dataset. For all the data, 70% was used for training, while 30% was reserved for testing. The model was provided with the data from the initial 30% of time points to predict the subsequent data points.

#### Handling Missing Observations

To handle missing observations during training, MicroProphet employs a self-supervised learning technique. During training, certain time points are randomly masked, and the model is required to predict these missing values based on the context provided by other available time points. This masking strategy enables the model to learn the inherent temporal patterns and dependencies within the time series even in the presence of missing observations, thus enhancing its robustness. Moreover, the introduction of positional encoding allows the model to treat missing observations as an implicit “blank” rather than simple zero-padding. Positional encoding enables the model to retain the relative position information of each time point, ensuring that even in the absence of certain data points, the model can still accurately capture the temporal dependencies within the sequence, maintaining the consistency of the predicted time series. Thus, MicroProphet is not only capable of making predictions in the presence of missing observations but also capable of learning and inferring the missing value, ultimately generating a complete time series. This capability allows MicroProphet to exhibit stronger adaptability and prediction accuracy when dealing with sparse or incomplete data.

#### SHAP Analysis for Microbial Feature Importance

To interpret the contribution of individual microbial taxa to time-series predictions, we employed SHapley Additive exPlanations (SHAP) analysis. The Shapley value ∅_*i*_ for a given feature *i* is defined as:

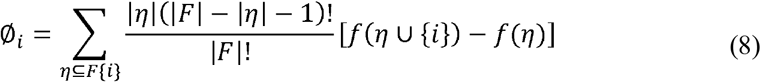

Where *F* is the set of all microbial taxa, *η* is a subset of *F* excluding feature *i*, *f(η*) is MicroProphet’s output when trained on subset *η*. This equation ensures that the contribution of each microbial taxon is fairly distributed across all possible feature sets, providing an interpretable importance score.

#### Evaluation Metrics

The evaluation of the model performance was conducted using both first-order metrics, which assess numerical accuracy, and second-order metrics, which evaluate ecological plausibility. First-order statistics include root mean squared error (RMSE), Pearson correlation and Cosine similarity, while the second-order metrics encompass indices such as the beta diversity index, among others.

### Degree of microbial individuality

The Degree of Microbial Individuality (DMI) quantifies the variability in microbial composition of a specific genus across individuals, with a higher DMI indicating greater individual-specific variation in microbial taxa. It is defined as the difference between the inter-individual and intra-individual Beta-diversity (BC distance) of a particular genus. The inter-individual BC distance reflects the variation in microbial composition between different individuals, while the intra-individual BC distance captures the variation within an individual over time. To compute DMI, we employ a bootstrap resampling technique^39^, which involves repeatedly resampling the data with replacement to assess the robustness and variability of the DMI estimates. Specifically, for each genus, the data are resampled and the DMI value is calculated using the following formula:

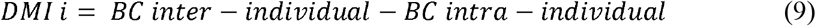

This resampling process is repeated multiple times to generate a distribution of DMI estimates, providing a comprehensive understanding of the stability and reliability of the DMI metric. In order to ensure the accuracy of the DMI estimates, we consider a DMI value reliable only if the standard deviation of the bootstrapped distribution is less than 15 times the mean DMI. This threshold helps to filter out any estimates that may be influenced by high variability or uncertainty in the data.

### Beta diversity analysis

The beta diversity was used as a measure of between-sample differences in community composition and was quantified as Bray-Curtis dissimilarity. The principal coordinate analysis (PCoA) was used to sort and visualize the beta diversity matrices. Differences in beta diversity between groups were tested using the ‘adonis2’ implementation of permutational multivariate analysis of variance (PERMANOVA).

### Local Similarity Network Analysis as a Directed and Weighted Networks

To uncover the temporal associations and interactions between microbial species, Local Similarity Analysis (LSA)^40^ was employed for network discovery. LSA is an advanced technique designed to capture both direct and time-lagged associations between species and environmental factors, without requiring significant data reduction. Traditional methods, such as Pearson Correlation Coefficient (PCC) analysis, are limited to identifying linear relationships and often fail to capture more complex, non-linear interactions that frequently occur in situ. By contrast, LSA excels at detecting these intricate dependencies, making it particularly useful for modeling ecological dynamics in microbial communities.

For two species-normalized time series *x*_i_(*t*) and *x*_j_ (*t*), we initialize two matrices: the positive score matrix *P*_*T* × *T*_ and the negative score matrix *N*_*T* × *T*_, both set to zero:

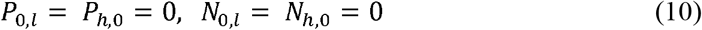

These matrices will store cumulative scores representing positive and negative correlations between *x*_*i*_*(t*) and *x*_*j*_*(t*) at different time points *t*_*l*_ and *t*_*h*_, *l*= 1,…,*T, h*= 1,…,*T*. For each time points *t*_*l*_ and *t*_*h*_, where |*t*_*l*_ − *t*_*h*_ | ≤ *D* (with *D* as a specified time lag threshold), the score matrices are updated iteratively.

The positive score matrix *P* is updated as follows:

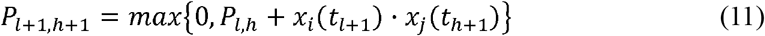

Similarly, the negative score matrix is updated as:

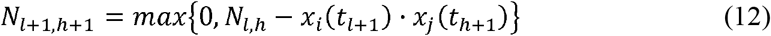

After iterating through all timepoints, the maximum positive and negative scores between species *i* and *j* are determined as follows:

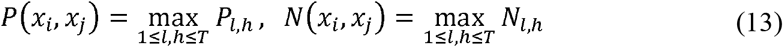

Then, the local similarity score *LS* (*x*_*i*_,*x*_*j*_) is computed by normalizing the maximum score by the total length of the time series *T*:

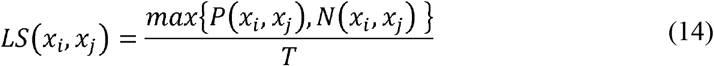

To determine whether the interaction is positive or negative, the sign of the similarity score is calculated as:

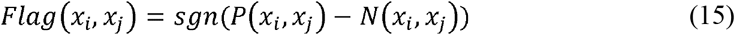

Last, the coefficient *A*_*ij*_ between species *i* and species *j* is obtained using local similarity score and its corresponding flag:

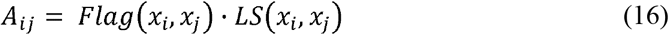

The application of LSA thus enhances the robustness and reliability of ecological network analysis, particularly when dealing with complex, interdependent biological systems.

To further determine the directional relationship between species, we applied the Kolmogorov-Smirnov (KS) test to compare the distribution of time series between two species. The KS test is a non-parametric statistical method used to evaluate the difference between two sample distributions. Specifically, the KS test calculates the maximum deviation between the two sample distributions and evaluates the significance of this difference using the following formula:

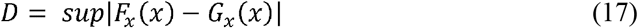

Where *F*_*x*_ (*x*) and *G*_*x*_(*x*) represent the cumulative distribution functions (CDFs) of species *x*_*i*_ and *x*_*j*_, respectively, and *D* is the KS test statistic.

By computing the CDFs of each species, we inferred the leading relationship between the species. If the CDF of species *x*_*i*_ precedes that of species *x*_*j*_, we consider species *x*_*i*_ as leading species *x*_*j*_, and vice versa. The directional relationship can be determined using the following condition:

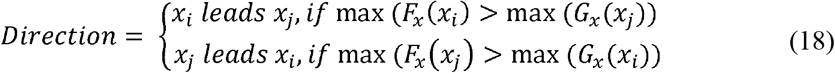

### UMAP for microbiome distribution

The distribution of the microbiome was visualized using Uniform Manifold Approximation and Projection (UMAP) with the R package “Seurat” (version 4.0). Prior to the application of UMAP, the count data were first normalized to relative abundance. A distance matrix was then computed using Bray-Curtis dissimilarity through the R package “Vegan” (version 2.6-2). Subsequently, Principal Coordinate Analysis (PCoA) was performed to transform the distance matrix. The UMAP projection was then calculated, and the results were visualized using the first two dimensions.

## Materials availability

This study did not generate new unique reagents.

## Data and code availability

All source code and data were available at https://github.com/HUST-NingKang-Lab/MicroProphet. Any additional information required to reanalyze the data reported in this paper is available from the lead contact upon request.

## Acknowledgements

This work was partially supported by the National Key R&D Program of China (Grant No. 2023YFA1800900 and 2018YFC0910502), the National Natural Science Foundation of China (Grant Nos. 32071465, 31871334, 81827901). Numerical computations were performed on the Hefei Advanced Computing Center. We also acknowledge BioRender (https://www.biorender.com/) for providing the tools used to create the illustrations in this work.

## Author information

### Contributions

KN conceived of and proposed the idea and designed the study. YZ and KZ designed and developed the framework. YZ, KZ, XC, HZ and JH performed the experiments and analyzed the data. YZ, KZ, and KN contributed to editing and proofreading the manuscript. All authors read and approved the final manuscript.

