## Supplementary figure for "MicroProphet: A Digital Twin Framework for Predicting Microbial Community Dynamics with Personalized Precision"

**Supplementary figures for**

**Processing time-series microbiome with MicroProphet digital twin framework improves microbial community dynamics prediction with personalized precision**

Yuli Zhang^1, #^, Kouyi Zhou^1, #^, Xiaoke Chen^1, #^, Haohong Zhang^1^, Jin Han^1^, Kang Ning^1, *^

1 Key Laboratory of Molecular Biophysics of the Ministry of Education, Hubei Key Laboratory of Bioinformatics and Molecular-imaging, Center of AI Biology, Department of Bioinformatics and Systems Biology, College of Life Science and Technology, Huazhong University of Science and Technology, Wuhan 430074, Hubei, China

^#^ These authors contributed equally to this work.

### **Supplementary Figure 1-7**

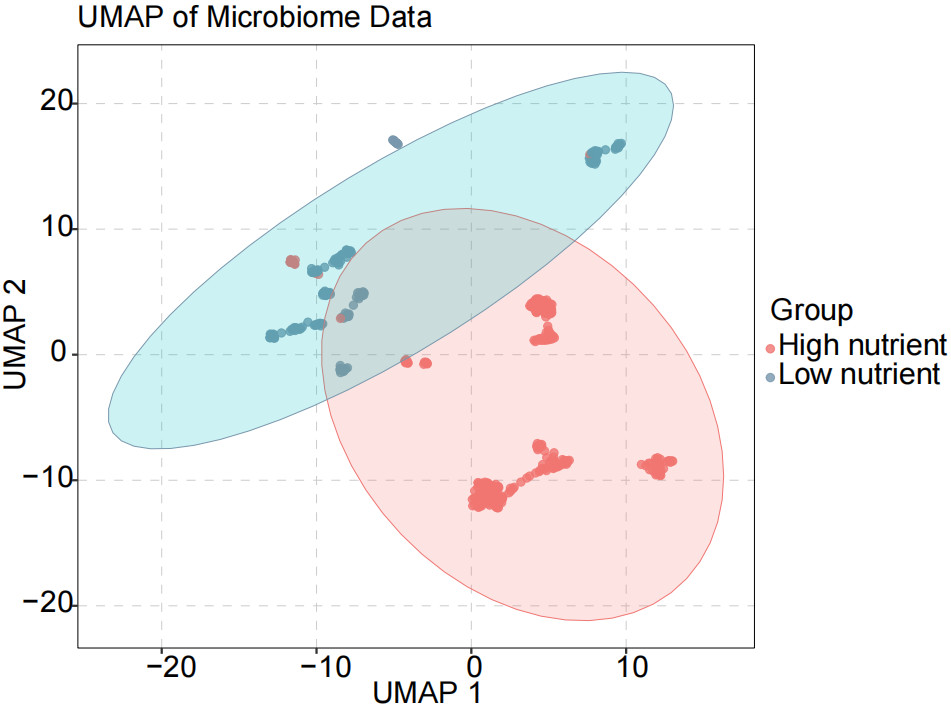

**Supplementary Figure 1. UMAP plot illustrating the microbial composition of synthetic data from high nutrient and low nutrient environments.** The colors represent the two groups, with red indicating high nutrient and blue indicating low nutrient. The low-nutrient condition consisting of 1 g l−1 of yeast extract, 1 g l−1 of soytone, 10 mM sodium phosphate, and trace elements, while the high-nutrient condition is supplemented with 5 g l−1 of glucose and 4 g l−1 of urea to increase interaction strength by amplifying resource competition and promoting environmental pH fluctuations.

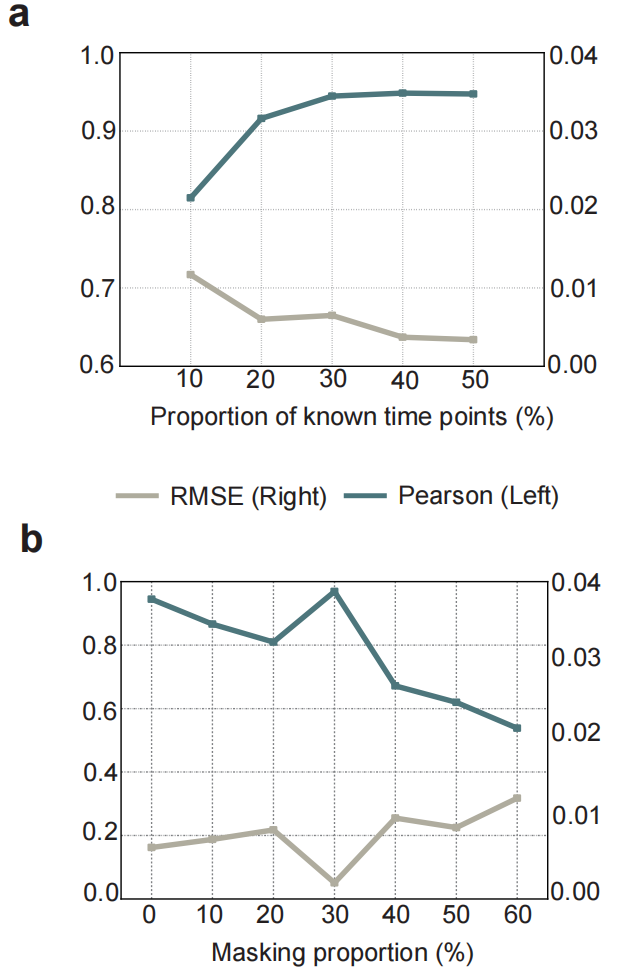

**Supplementary Figure 2. Performance evaluation of prediction models using varying time points and data masking proportions.** The Pearson correlation coefficient is shown with a dark blue line, while the RMSE is represented by the light gray line.

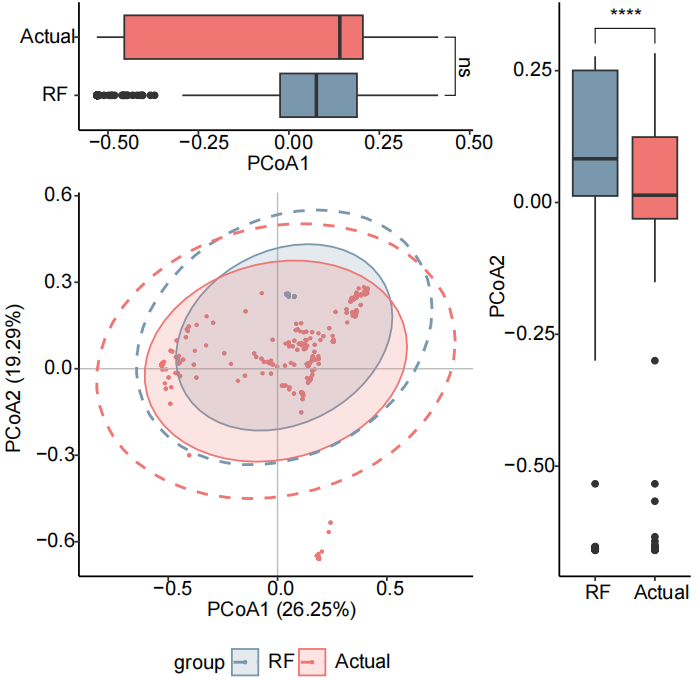

**Supplementary Figure 3. Beta diversity analysis before and after interpolation using Random Forest.** The blue color represents predicted values, while the red color represents actual values.

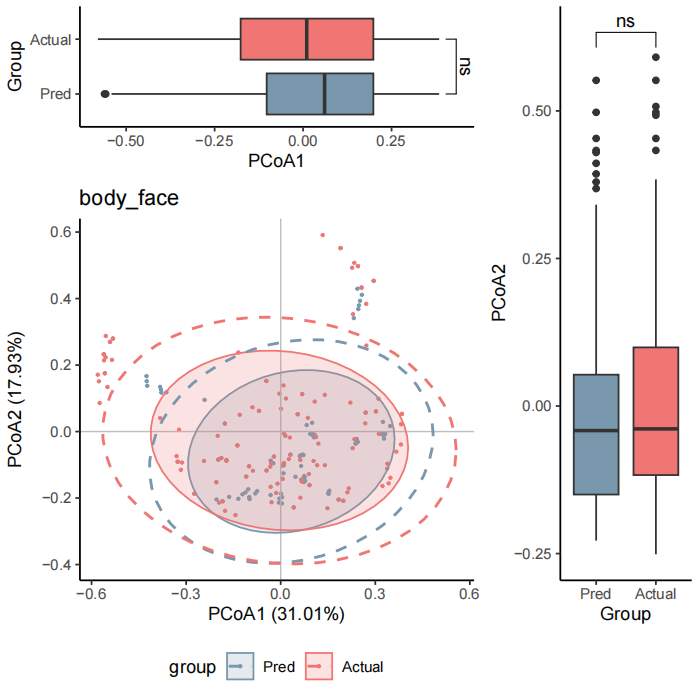

**Supplementary Figure 4. Beta diversity analysis of IBD dataset comparing predicted and actual dataset.** The blue color represents the data predicted by MicroProphet, while the red color represents the actual observational dataset.

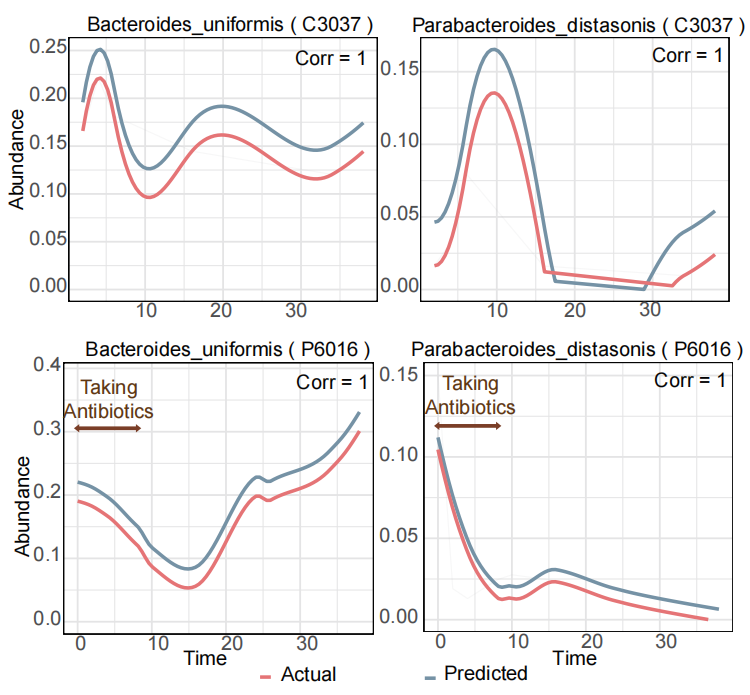

**Supplementary Figure 5. Fitting of microbial abundance trajectories under antibiotic intervention.** The top and bottom panels illustrate distinct temporal abundance trends for the same taxa across two subjects: C3037, who did not take antibiotics, and P6016, who underwent antibiotic treatment. The red lines represent the actual abundances, while the blue lines represent the predicted abundances. Shaded areas indicate confidence intervals for the predictions. The horizontal axis represents time, and the vertical axis represents microbial abundance.

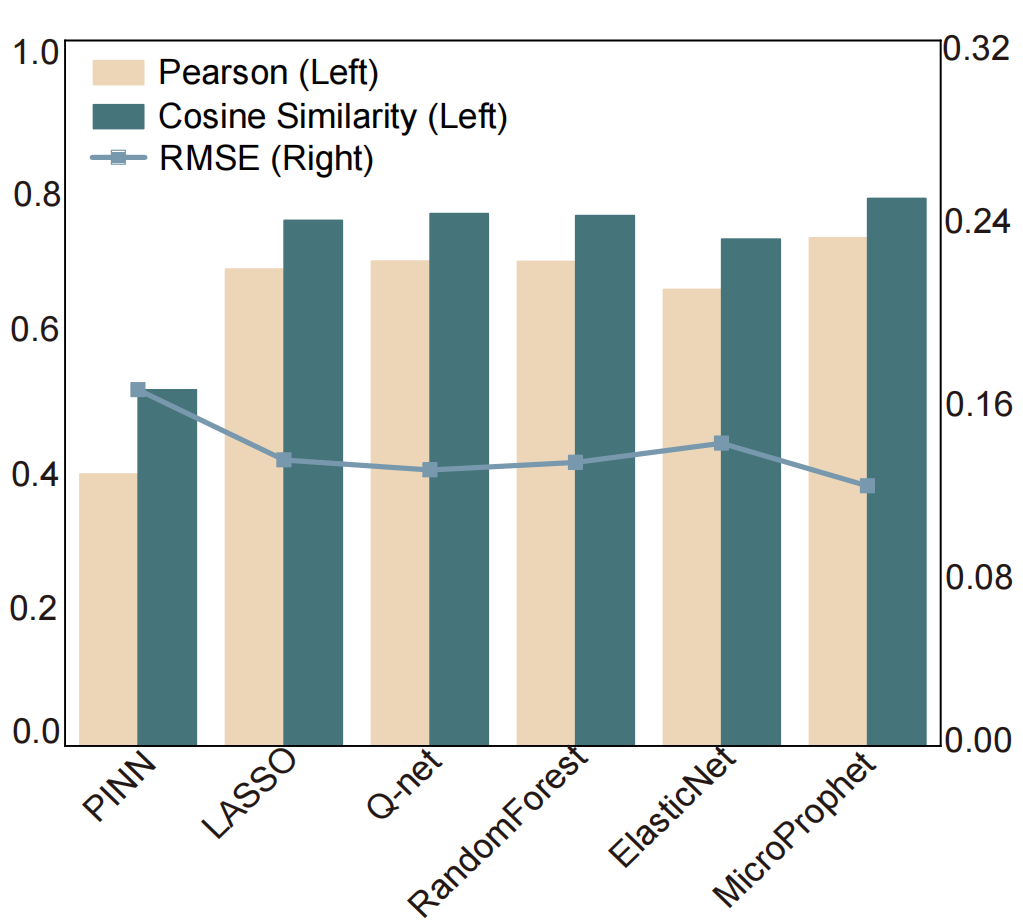

**Supplementary Figure 6. Comparison of prediction performance for different methods using the Corpse dataset.** The left panel shows Pearson and Cosine Similarity metrics (in yellow and green respectively), while the right panel shows RMSE (in blue).

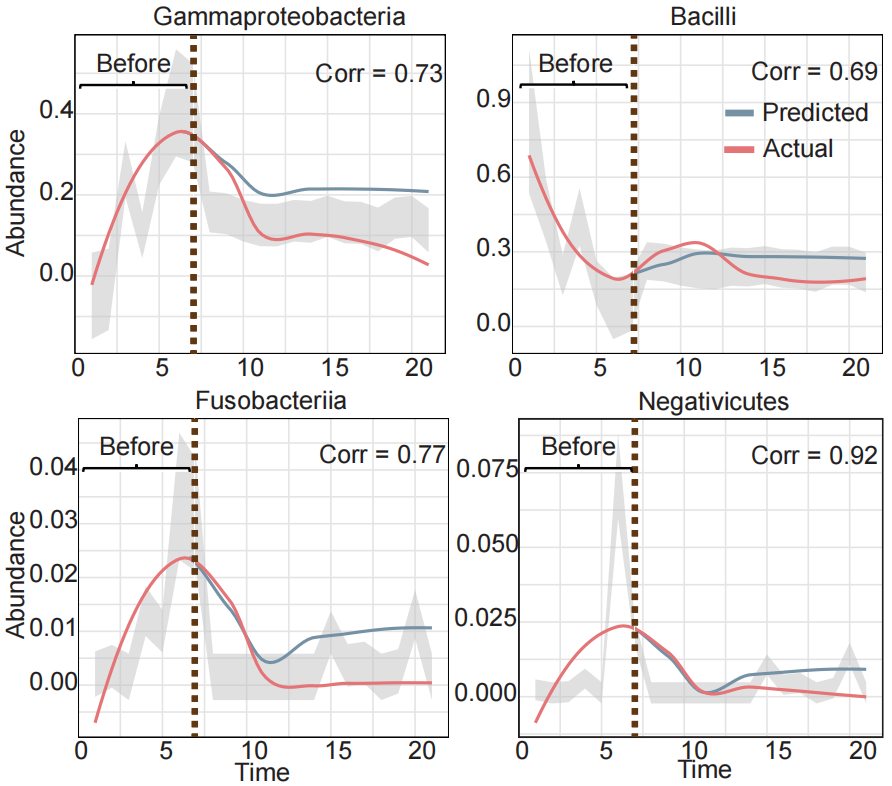

**Supplementary Figure 7. Predicted trajectories of highly abundant taxa in Corpse dataset.** Line plots display the temporal abundance trajectories of four representative microbial taxa (Gammaproteobacteria, Bacilli, Fusobacteria, and Negativicutes) over the 21 days of decomposition. The red lines represent the actual abundances, while the blue lines represent the predicted abundances. Shaded areas indicate confidence intervals for the predictions. The horizontal axis represents time, and the vertical axis represents microbial abundance.
